# Functional Organization of Mouse Primary Auditory Cortex in adult C57BL/6 and F1 (CBAxC57) mice

**DOI:** 10.1101/843300

**Authors:** Zac Bowen, Daniel E. Winkowski, Patrick O. Kanold

## Abstract

The primary auditory cortex (A1) plays a key role for sound perception since it represents one of the first cortical processing stations for sounds. Recent studies have shown that on the cellular level the frequency organization of A1 is more heterogeneous than previously appreciated. However, many of these studies were performed in mice on the C57BL/6 background which develop high frequency hearing loss with age making them a less optimal choice for auditory research. In contrast, mice on the CBA background retain better hearing sensitivity in old age. Since potential strain differences could exist in A1 organization between strains, we performed comparative analysis of neuronal populations in A1 of adult (~10 weeks) C57BL/6 mice and CBAxC57 F1 mice. We used *in vivo* 2-photon imaging of pyramidal neurons in cortical layers L4 and L2/3 of awake mouse primary auditory cortex (A1) to characterize the populations of neurons that were active to tonal stimuli. Pure tones recruited neurons of widely ranging frequency preference in both layers and strains with neurons in CBA mice exhibiting a wider range of frequency preference particularly to higher frequencies. Frequency selectivity was slightly higher in C57BL/6 mice while neurons in CBA mice showed a greater sound-level sensitivity. The spatial heterogeneity of frequency preference was present in both strains with CBA mice exhibiting higher tuning diversity across all measured length scales. Our results demonstrate that the tone evoked responses and frequency representation in A1 of adult C57BL/6 and CBAxC57 F1 mice is largely similar.

## Introduction

The cerebral cortex is uniquely adapted to encode behaviorally relevant stimuli and generate appropriate behavioral actions. The primary auditory cortex (A1) plays a key role for sound perception since it represents one of the first cortical processing stations for sounds. Recent studies have investigated the functional organization and circuitry of mouse auditory cortex (Bandyopadhyay et al., 2010; Rothschild et al., 2010; Li et al., 2017; Meng et al., 2017; Liu et al., 2019; Tischbirek et al., 2019). A classic hallmark of A1 organization in all species has been the discovery of tonotopic maps which describe a smooth distribution of tone preference across the surface of the primary and secondary auditory fields when probed with low spatial resolution techniques (Merzenich et al., 1975; Reale and Imig, 1980; Stiebler et al., 1997; Wessinger et al., 1997; Nelken et al., 2004; Bizley et al., 2005; Woods et al., 2010; Striem-Amit et al., 2011; van Dijk and Langers, 2013; Issa et al., 2014; Moerel et al., 2014; Baba et al., 2016; Liu et al., 2019). However, the application of large-scale high-resolution optical techniques in mice or rats using synthetic dyes (OGB1, Fluo4, or Cal-520) or sensitive genetically encoded Ca^2+^ indicators (GCaMP6) has shown that on the cellular level the organization of tuning in A1 is more heterogeneous than previously appreciated (Bandyopadhyay et al., 2010; Rothschild et al., 2010; Winkowski and Kanold, 2013; Kanold et al., 2014; Li et al., 2017; Meng et al., 2017; Panniello et al., 2018; Liu et al., 2019; Tischbirek et al., 2019; Zeng et al., 2019) suggesting that at least in rodent layer 2/3 (L2/3) functional tonotopic maps might be fractured. The precise degree of heterogeneity varies slightly between these studies, and this difference might be due to differences in anesthetic condition, cell inclusion criteria, tonal stimuli used, and/or species.

Many functional studies in mouse have been performed on mice on the C57BL/6 (C57) background. However, the C57 strain carries a mutant *cdh23* allele (Di Palma et al., 2001; Zheng et al., 2005; Zheng et al., 2009; Johnson et al., 2010; Kane et al., 2012; Miyasaka et al., 2013; Mock et al., 2016) which leads to a progressive loss of hair cells from the basal turn of the cochlea (Park et al., 2010), making these mice an ideal model of presbycusis (Hunter and Willott, 1987; Parham and Willott, 1988; Li and Borg, 1991; Willott et al., 1994; Ouagazzal et al., 2006). In contrast, mice of the CBA strain or F1 offspring of C57 mice and CBA mice retain normal hearing into adulthood (Hunter and Willott, 1987; Parham and Willott, 1988; Li and Borg, 1991; Willott et al., 1994; Ouagazzal et al., 2006; Frisina et al., 2011). Thus, potential hearing loss present at the ages studied in C57 mice could have contributed to the observation of local frequency preference heterogeneity. We aimed to test if the laminar differences in functional responses and organization present under anesthetized conditions in C57 mice were also present in awake animals and if such organizational features were also present in mice on the CBA background.

We performed *in vivo* 2-photon imaging experiments in awake mice of C57 strain as well as CBA background (CBAxC57 F1) and quantified the functional responses across L2/3 and L4 of A1. We focused on young adult animals (~2-3 months old) which is the age frequently used in functional studies, especially those involving mouse behavior (Bandyopadhyay et al., 2010; Rothschild et al., 2010; Carcea et al., 2017; Guo et al., 2017; Kuchibhotla et al., 2017; Li et al., 2017; Meng et al., 2017; Resnik and Polley, 2017; Francis et al., 2018; Liu et al., 2019; Tischbirek et al., 2019). To compare strains we imaged the mid-frequency region of A1 (4-16kHz) which is also where mice are most sensitive to sound (Zheng et al., 1999). We find that A1 of CBA mice contains more neurons tuned to high frequencies and a higher number of significant tone-responses overall, including a higher proportion of off-responses (Liu et al., 2019). Moreover, on the single cell level, bandwidth was slightly lower in L2/3 of C57 mice while rate-level curves revealed an increased gain in CBA mice. On the population level, we find that in both strains the local heterogeneity is higher in L2/3 than in L4 but that heterogeneity is slightly higher in CBA mice than C57 mice. Our observations show that while there are differences in the tuning of single neurons between mice, there remains laminar differences in both frequency selectivity and local heterogeneity of frequency preference present in both strains. Our results here along with previously published results show that the local heterogeneity of frequency preference is present in both anesthetized and awake mice and across mouse strains. Thus, the heterogeneity of pure tone frequency selectivity forms an organizational principle of auditory cortex organization in mice.

## Methods

All animal procedures and experiments were approved by the University of Maryland Institutional Animal Care and Use Committee to be in accordance with applicable guidelines and regulations.

Mice were housed under a reversed 12 h-light/12 h-dark light cycle with ad libitum access to food and water. Imaging experiments were generally performed near the end of the light and beginning of the dark cycle. We imaged adult mice (C57BL/6: P46-P100; CBAxC57 F1: P66-P178 at time of imaging). Mice of both sexes were used based on their availability not by any biased selection.

### Animals

Mice used for this study expressed the genetically encoded calcium indicator GCaMP6s (Chen et al., 2013). We used transgenic mice that expressed GCaMP6s under the Thy1 promoter (Jax: 024275, GP4.3) conditionally when crossed to a mouse driver line expressing Cre recombinase under the control of Emx1 (GCaMP6s mouse: Jax: 024115; Emx1-Cre mouse: Jax: 005628). Mice on this background do not show epileptiform activity in A1 (Bowen et al., 2019). Mice on the C57BL/6 background show early onset age related hearing loss (Hunter and Willott, 1987; Parham and Willott, 1988; Li and Borg, 1991; Willott et al., 1994; Ouagazzal et al., 2006; Frisina et al., 2011). To compare these mice with mice that do not develop early hearing loss hearing we used the F1 generation of a hybrid mouse line, CBA/CaJ (Jax: 000654) crossed with C57BL/6-Thy1 which have normal hearing (Frisina et al., 2011). L2/3 and L4 datasets were always obtained sequentially in the same mouse, meaning the experiment (sound presentations) was conducted in one layer and then sequentially conducted in the other layer. The two transgenic lines exhibited robust indicator expression in both L2/3 and L4 (Bowen et al., 2019). We found no significant differences of indicator expression within layers when we compared across transgenic mouse lines and animals with viral expression and thus combined the data from different sources according to laminar position.

### Surgery and animal preparation

Mice were given a subcutaneous injection of dexamethasone (5mg/kg) at least 2 hours prior to surgery to reduce potential inflammation and edema from surgery. Mice were deeply anesthetized using isoflurane (5% induction, 2% for maintenance) and given subcutaneous injections of atropine (0.2 mg/kg) and cefazolin (500 mg/kg). Internal body temperature was maintained at 37.5 °C using a feedback-controlled heating blanket. The scalp fur was trimmed using scissors and any remaining fur was removed using Nair. The scalp was disinfected with alternating swabs of 70% ethanol and betadine. A patch of skin was removed, the underlying bone was cleared of connective tissue using a bone curette, the temporal muscle was detached from the skull and pushed aside, and the skull was thoroughly cleaned and dried. A thin layer of cyanoacrylate glue (VetBond) adhesive was applied to the exposed skull surface and a custom machined titanium head plate (based on the design described in Guo et al. (2014) was affixed to the skull overlying the auditory cortex using VetBond followed by dental acrylic (C&B Metabond). A circular craniotomy (~3 mm diameter) was made in the center opening of the head plate and the patch of bone was removed. Then, a chronic imaging window was implanted. The window consisted of a stack of 2 – 3 mm diameter coverslips glued with optical adhesive (Norland 71, Edmund Optics) to a 5 mm diameter coverslip. The edges of the window between the glass and the skull were sealed with a silicone elastomer (Kwik-Sil) and then covered with dental acrylic. The entire implant except for the imaging window was then coated with black dental cement created by mixing standard white powder (Dentsply) with iron oxide powder (AlphaChemical, 3:1 ratio) (Goldey et al., 2014). Meloxicam (0.5 mg/kg) and a supplemental dose of dexamethasone were provided subcutaneously as a post-operative analgesic. Animals were allowed to recover for at least 1 week prior to the beginning of experiments.

### Acoustic stimulation

Sound stimuli were synthesized in MATLAB using custom software (courtesy of P. Watkins, UMD), passed through a multifunction processor (RX6, TDT), attenuated (PA5, Programmable Attenuator), and delivered via ES1 speaker placed ~5 cm directly in front of the mouse. The sound system was calibrated between 2.5 and 80 kHz and showed a flat (±3 dB) spectrum over this range. Overall sound pressure level (SPL) at 0 dB attenuation was ~90 dB SPL (for tones). Sounds were played at three sound levels (40, 60, and 80 dB SPL). Auditory stimuli consisted of sinusoidal amplitude-modulated (SAM) tones (20 Hz modulation, cosine phase), ranging from 3 – 48 kHz. For wide-field imaging, the frequency resolution of the stimuli was 1 tone/octave; for 2-photon imaging, the frequency resolution was 2 tones/octave (0.5 octave spacing). Each of these tonal stimuli was repeated 5 times with a 4 – 6 s interstimulus interval for a total of either 75 (wide-field) or 135 (2-photon) iterations.

### 2-photon imaging

To study cellular neuronal activity using 2-photon imaging, we used a scanning microscope (Bergamo II series, B248, Thorlabs) coupled to a pulsed femtosecond Ti:Sapphire 2-photon laser with dispersion compensation (Vision S, Coherent). The microscope was controlled by ThorImageLS software. The laser was tuned to a wavelength of λ = 940 nm in order to simultaneously excite GCaMP6s and mRuby2. Red and green signals were collected through a 16× 0.8 NA microscope objective (Nikon). Emitted photons were directed through 525/50-25 (green) and 607/70-25 (red) band pass filters onto GaAsP photomultiplier tubes. The field of view was 370×370 μm^2^. Imaging frames of 512×512 pixels (0.52 μm^2^ pixel size) were acquired at 30 Hz by bi-directional scanning of an 8 kHz resonant scanner. Beam turnarounds at the edges of the image were blanked with a Pockels cell. The average power for imaging in both L2/3 and L4 was <~70 mW, measured at the sample plane. We imaged in the mid-frequency regions of A1 and validated this by measuring the median BF of cells in the imaging field.

### Data Analysis

Wide-field imaging was used to identify the large scale tonotopic organization as previously (Meng et al., 2017; Liu et al., 2019). Wide-field image sequences were analyzed using custom routines written in Matlab (Mathworks). Images were parsed into trial-based epochs in which each frame sequence represented a single trial consisting of the presentation of a single sound frequency-intensity combination. For each trial, response amplitude (ΔF/F_0_) as a function of time was determined for each pixel using the formula ((F – F_0_)/ F_0_) where F corresponds to the time varying fluorescence signal at a given pixel and F_0_ was estimated by averaging the fluorescence values over 4 frames (~1 s) prior to sound onset for a given trial and pixel. For construction of sound-evoked response maps, the amplitude of the ΔF/F_0_ pixel response during 1 s after stimulus onset (~4 frames) was averaged across time and repetitions yielding an average response magnitude that was assigned to each pixel. Responsive areas in the average response maps were defined on a pixel-by-pixel basis as pixels in which the average brightness of the pixel during the 1 s after stimulus onset exceeded 2 standard deviations of the pixel brightness during the 1 s before the stimulus across stimulus repetitions.

### 2-photon image analysis

Image sequences were corrected for x-y drifts and movement artifacts using either the TurboReg in ImageJ (Thevenaz et al., 1998; Schindelin et al., 2012) or discrete Fourier transform registration (Guizar-Sicairos et al., 2008) implemented in Matlab (Mathworks). Neurons were identified manually from the average image of the motion corrected sequence. Ring-like regions of interest (ROI) boundaries were drawn based on the method described in Chen et al. (Chen et al., 2013). Overlapping ROI pixels (due to closely juxtaposed neurons) were excluded from analysis. For each labeled neuron, a raw fluorescence signal over time (F) was extracted from the ROI overlying the soma. The mean fluorescence for each neuron was calculated across frames and converted to a relative fluorescence measure (ΔF/F_0_), where ΔF = (F – F_0_). F_0_ was estimated by using a sliding window that calculated the average fluorescence of points less than the 50^th^ percentile during the previous 10-second window (300 frames). Neuropil (NP) subtraction was performed on all soma ROIs (Peron et al., 2015). In short, the neuropil ROI was drawn based on the outer boundary of the soma ROI and extended from 1 pixel beyond the soma ROI outer boundary to 15 μm excluding any pixels assigned to neighboring somata. Thus, the final ΔF/F_0_ used for analysis was calculated as ΔF/F_0_ = (ΔF/F_0_)_soma_ – (α x (ΔF/F_0_)_NP_), where we used α = 0.9 (Peron et al., 2015) to reduce fluorescence contamination from the neuropil. Neurons in which the ΔF/F_0_ signal was significantly modulated by sound presentation were defined by ANOVA (p < 0.01) across baseline (pre-stimulus) and all sound presentation periods. Single neuron receptive fields (RF) were determined as the average ΔF/F_0_ response to each frequency-intensity combination across 5 stimulus repetitions during the stimulus window (1 s). Best frequency (BF) for each neuron was determined as the center of mass of the RF. The characteristic frequency (CF) is the highest average dF/F response at the lowest sound level for which the neuron had a significant response.

Receptive Field Sum (RFS) was calculated by first only keeping values of the RF for which the neuron had a significant response above baseline (setting non-significant elements to zero). This RF was then normalized and summed across all elements to arrive at the RFS (Fig. 2A). For a perfectly selective neuron that only responded significantly to one frequency/SPL combination, the RFS would equal 1. The closer the RFS was to 1, the more selective the neuron. A higher RFS indicates a neuron that was highly responsive across a large area of its receptive field. The theoretical maximum value would be a neuron responding significantly and equally to all frequency/SPL combinations giving a value of 36, however this was never observed in our data.

## Results

To investigate the functional responses of A1 neurons and their functional organization, we performed in vivo 2-photon imaging in awake animals chronically implanted with cranial windows (Meng et al., 2017; Liu et al., 2019) using GCaMP6s expression. As done previously we performed widefield imaging to localize the imaging fields to A1 (Meng et al., 2017; Liu et al., 2019). We acquired L2/3 and L4 data sets sequentially in the same mouse ensuring that we imaged in the same tonotopic position in the mid-frequency region (Fig. 1A). We imaged 22 fields of view (FOV) from C57BL/6 (C57) mice (11 L2/3, 11 L4) and 35 FOVs in CBA mice (18 L2/3, 17 L4). Imaging depths in C57 mice were 177±17 µm (mean±std) for L2/3 and 415±7 µm for L4 and in CBA mice 160±17 µm for L2/3 and 412±5 µm for L4.

**Figure 1.**
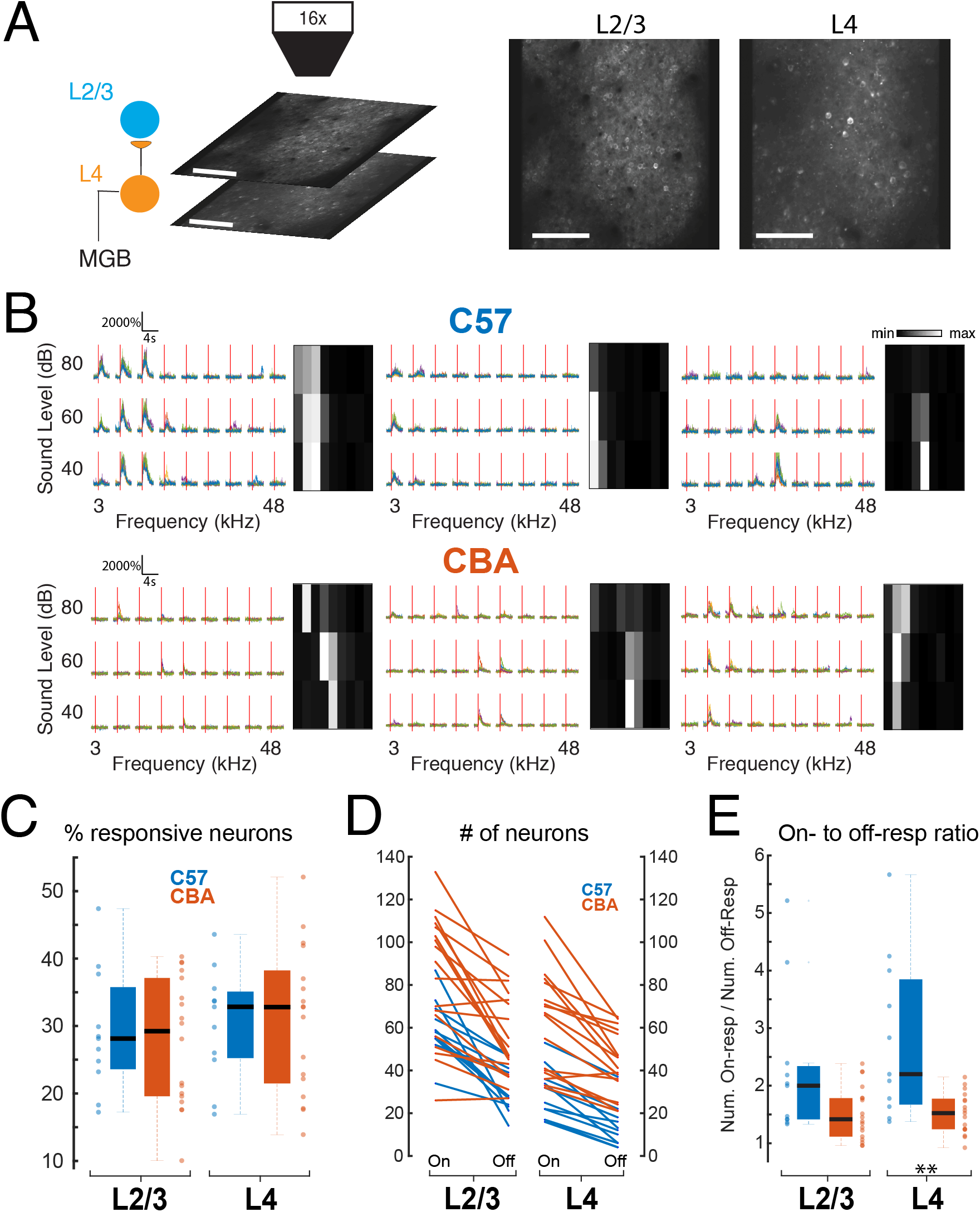
Neurons respond to a range of stimuli in both layers of both strains. **A)** Example 2-photon imaging FOVs in L2/3 and L4 of awake mouse auditory cortex. Scale bar 100µm. **B)** Example calcium traces and frequency response areas (Receptive Fields) for C57 mice (top) and CBA mice (bottom). Red vertical lines indicate tone-onset. Individual traces represent single trial calcium transients. **C)** Percent of neurons in each FOV deemed significantly responding to at least one stimulus. Blue indicates C57 mice and orange indicates CBA mice. **D)** Number of neurons deemed On-responsive and Off-responsive in each FOV. Lines connect number of cells from the same FOV. **E)** Ratio of on-to off-responsive neurons for each of the FOVs plotted in **D**. ** indicates significance at p<0.01 (Two-sample t-test).

### A similar fraction of neurons responded to tonal stimuli across strains with respect to cortical layer

We first investigated the fraction of single neurons in L4 and L2/3 of A1 in mice of both strains that respond to tonal stimuli. We played a range of tones (3-48 kHz, half-octave spacing) at 4 sound pressure levels (80, 60, 40, and 20 dB) with 5 repetitions of each stimulus and found a wide variety of receptive fields, or frequency response areas (FRA), across the recorded neuronal populations (Fig. 1B). We identified tone-responsive neurons by a significant increase in the somatic fluorescent signal above baseline (dF/F) during stimulus presentation. We found that the proportion of neurons in each FOV with significant responses to at least one stimulus were similar (~25-35%) in both mouse strains and across cortical layers (Fig. 1C).

While many A1 neurons have a maximal response during the stimulus, A1 neurons can also have offset responses to tones which can be separated from each other with long (>1s) stimuli (Liu et al., 2019). We measured how many neurons had significant onset (tone-on) responses versus offset (tone-off) responses (Fig. 1D). We found that both CBA and C57 mice had tone-on as well as tone-off neurons with more tone-on neurons being present than tone-off neurons in both strains. Overall, L4 datasets had fewer lower number of neurons responding. We computed an on/off ratio as the ratio between the number of on-responsive neurons to the number of off-responsive neurons to quantify the disparity between on- and off-responsive neurons in each FOV. We found that the ratio is higher in both L2/3 and L4 of C57 mice than CBA mice (Fig. 1E) (p_L2/3_<0.06, Wilcoxon rank sum, p_L4_<0.002, two-sample t-test). Together, these results indicate that C57 mice have fewer off-responsive neurons relative to the number of on-responsive neurons, particularly in L4 than CBA mice. These results indicate that, in general, A1 of both strains contains a similar number of tonally responsive neurons but that L4 of CBA mice contains significantly more off-responsive neurons when compared to C57 mice.

### A1 neurons in CBA mice have a broader range of tuning than C57 mice

Having found that overall neuronal responsiveness was similar between strains, we next investigated the tuning properties of single neurons in both mouse strains. C57 mice show a progressive loss of high frequency hearing (Hunter and Willott, 1987; Parham and Willott, 1988; Li and Borg, 1991; Willott et al., 1994; Ouagazzal et al., 2006; Park et al., 2010). We thus investigated the possibility that A1 neurons in C57 mice showed altered frequency preference. Moreover, at C57 mice show a loss of parvalbumin expression with age suggesting a decrease in inhibition (Martin del Campo et al., 2012). Since inhibition sharpens receptive fields, we also investigated the possibility that A1 neurons in C57 mice showed altered frequency selectivity.

We first investigated the frequency preference of neurons by calculating the best frequency (BF) and characteristic frequency (CF) of each significantly responding neuron. The BF is the frequency that produced the highest response in each neuron regardless of sound pressure level, whereas the CF is the frequency that produced the highest response at the lowest sound pressure level that elicited a significant response (Fig. 2A).

**Figure 2.**
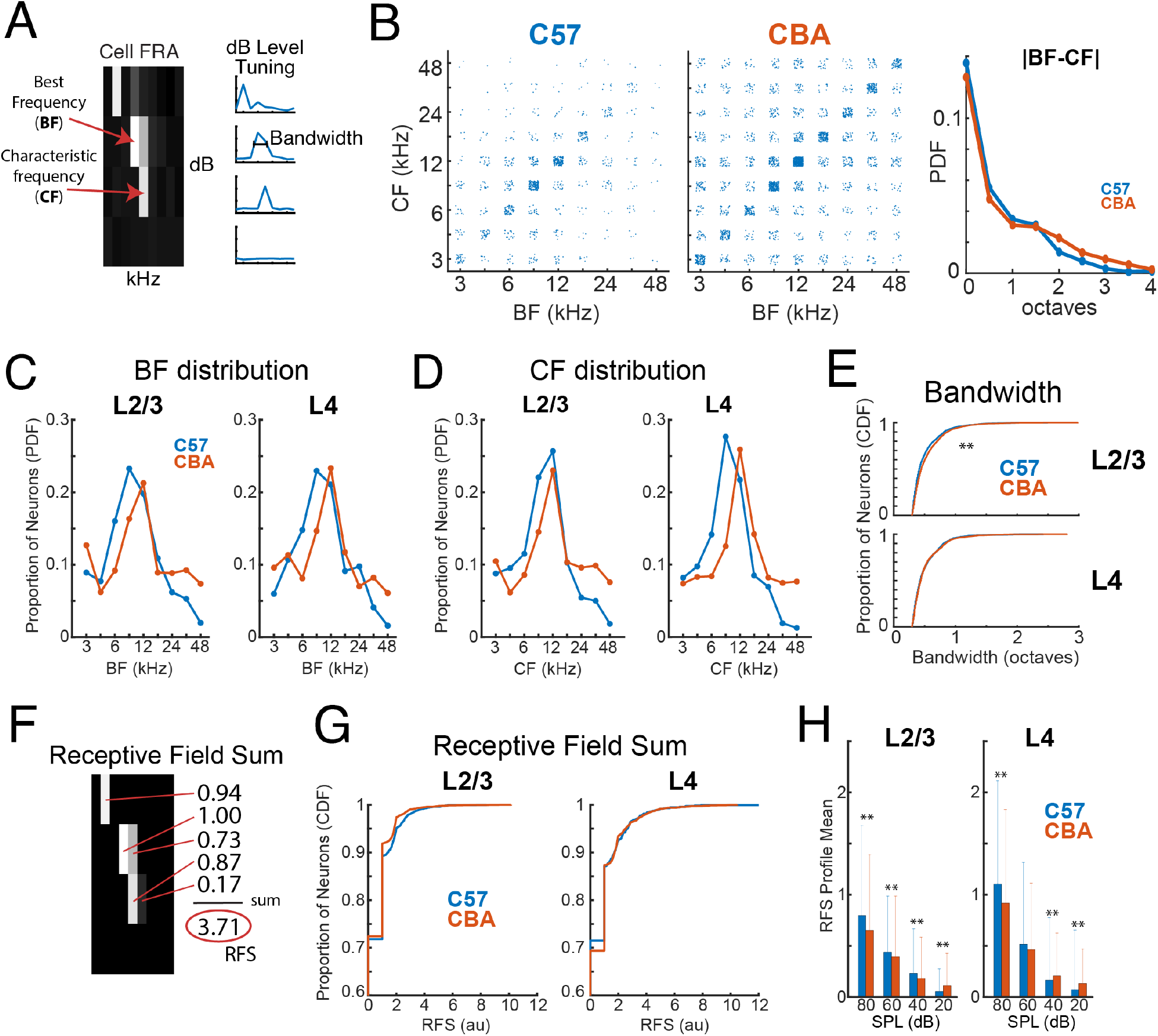
Similar tuning and selectivity across mouse strains. **A)** Example FRA illustrating the measurement and computation of BF, CF, and bandwidth. **B)** BF plotted against CF for each neuron in both mouse strains. Each dot represents an individual neuron. |BF-CF| shows the absolute magnitude of difference between BF and CF for each mouse strain. **C)** Histograms of BF distributions for C57 (blue) and CBA (orange) mouse strains in both L2/3 (left) and L4 (right). **D)** Conventions as in **C**, but for CF. **E)** CDFs of bandwidth in C57 (blue) and CBA (orange) mouse strains in both L2/3 (left) and L4 (right). ** indicates significance at p<0.01 (Wilcoxon rank sum test). **F)** Example calculation of Receptive Field Sum. **G)** CDFs of receptive field sum values in C57 (blue) and CBA (orange) mouse strains in both L2/3 (left) and L4 (right). **H)** Receptive field sum split up by sound level. Sum is taken across rows in **F** and then averaged across all neurons. ** indicates significance at p<0.01 (Wilcoxon rank sum test).

We find that the BF and CF distributions are similar across L2/3 and L4 within each strain of mice (Fig. 2C,D). However, comparing strains showed that C57 mice had a somewhat broader distribution of represented BFs in the mid-frequency region (6-17kHz) but a reduced number of neurons representing the high end of the frequency range (33-48kHz) (Fig. 2C,D). These data indicate that the representation of tonal stimuli differs between mouse strains and that C57 mice show an under-representation of high-frequency stimuli consistent with high frequency hearing loss.

BF and CF measure different aspects of the FRA, and the BF can be affected by non-linear rate-level changes at the CF. If all single neurons have a linear increase in responses with increasing sound level, then CF and BF would be equal. Thus, we next probed the relationship between CF and BF for individual cells by plotting BF versus CF for each neuron in both mouse strains (combined across layers) (Fig. 2B). To account for differences in number of points, we also plot the PDF of |BF-CF| across all neurons (Fig. 2B, ***right***). We find that while CF and BF are mostly correlated in both mouse strains, C57 mice have fewer neurons with a mismatched BF and CF. This may be a consequence of C57 mice having worse detection of quieter sounds (lower SPL). As a result, each neuron may only have significant responses at one or two SPLs causing the maximal FRA response (BF) to occur at the lowest SPL with a significant response (CF). Moreover, the slightly broader BF vs CF distribution in CBA mice could also indicate that cells respond to a broad range of frequencies at varying SPLs, whereas cells in C57s mice are often sharply tuned.

We next assessed how frequency-selective A1 neurons were by calculating the bandwidth of the tuning curve at 70% of BF amplitude at the SPL for which BF was measured. Overall, C57 mice had narrower bandwidths than the CBA mice in L2/3, but the effect size is modest. These results indicate that C57 mice had a higher selectivity for tonal stimuli (Fig. 2E, ***left***). In contrast, bandwidth distributions between L4 from both mouse strains were similar (Fig. 2E, ***right***).

A1 responses can have nonlinear changes in magnitude with increasing sound level, thus bandwidth measures might miss responses outside traditional “V-shaped” FRAs. Thus, to further investigate the frequency selectivity of each neuron, we developed a measure called the Receptive Field Sum (RFS) which is more inclusive when quantifying nonlinear tuning curves or receptive fields. For each neuron this measure is calculated by first only keeping values of the receptive field (RF) for which the neuron had a significant response above baseline (setting non-significant elements to zero). The remaining RF elements are then normalized and summed to arrive at the RFS (Fig. 2F). A lower RFS indicates that a neuron was more specific in its stimulus responsiveness. We find that in L2/3 neurons, CBA mice have a slightly lower distribution of RFS values than C57 mice (Fig. 2G, ***left***). Since C57 L2/3 populations have narrower bandwidth (Fig. 2E, ***left***), the higher RFS may indicate that C57 mice have a higher magnitude response to a select few stimuli in their receptive field consistent with reduced inhibition (Martin del Campo et al., 2012). However, L4 neurons did not differ in bandwidth or RFS values between strains (Fig. 2G, ***right***), indicating that any changes in receptive fields as a result of age-related hearing loss is only reflected in layer 2/3. To investigate further, we split up RFS by sound level by summing the normalized receptive field across frequencies, rather than across both frequencies and sound levels (Fig. 2H). This gives a SPL-dependent profile of significant responses and their magnitude with respect to the rest of the RF. We find that in L2/3, C57 mice have a significantly higher RFS profile mean except at 20 dB where CBA is higher. In L4 the CBA mice have a significantly lower RFS profile mean at 80 dB yet is significantly higher at 40 dB and 20 dB. From these results we observe that overall C57 mice had a much more drastic decrease in significant responses at quieter sound levels than in CBA mice. We conclude that CBA mice retain better hearing sensitivity to quieter sounds in old age.

Together, these analyses show that mid-frequency regions of A1 of C57 mice contain frequency selective neurons with a frequency preference that is shifted towards lower frequencies than that of CBA mice. However, the frequency selectivity and responsiveness of individual neurons from C57 mice is slightly higher than that of cells in CBA mice particularly in L2/3.

### CBA mice have a higher local heterogeneity of frequency preference than C57 mice

While on large scales A1 in rodents shows a tonotopic organization, nearby neurons can show different frequency preference (Bandyopadhyay et al., 2010; Rothschild et al., 2010; Winkowski and Kanold, 2013; Kanold et al., 2014; Li et al., 2017; Meng et al., 2017; Panniello et al., 2018; Liu et al., 2019; Tischbirek et al., 2019; Zeng et al., 2019). We thus investigated if this local heterogeneity varied between mouse strains. We computed median, standard deviation (STD), and interquartile range (IQR) of the best frequency (BF) distribution for each FOV, where BF is the frequency that produced the highest response in each neuron regardless of sound pressure level. While IQR has been frequently used to assess ACX tuning heterogeneity, we further include the STD in order to include every observed tuned neuron in the field of view (i.e. even outliers). Consistent with the higher prevalence of high-frequency neurons in CBA mice (Fig. 2), the median BF of C57 FOVs in both L2/3 and L4 was 8.5 kHz while CBA mice has a median BF of 12 kHz in both layers (Fig. 3A). The IQR_BF_ in each FOV was slightly higher in CBA mice than C57 mice in both L2/3 and L4 (Fig. 3B). Despite the lack of significant difference of IQR_BF_ between strains, we observed significantly higher STD_BF_ of CBA mice in both L2/3 and L4 (Fig. 3C) (p_L2/3_<0.005, p_L4_<0.004, Wilcoxon rank sum and two-sample t-test respectively), indicating a strong presence of neurons tuned to vastly different frequencies than the rest of the FOV (i.e. outliers) in CBA mice.

**Figure 3.**
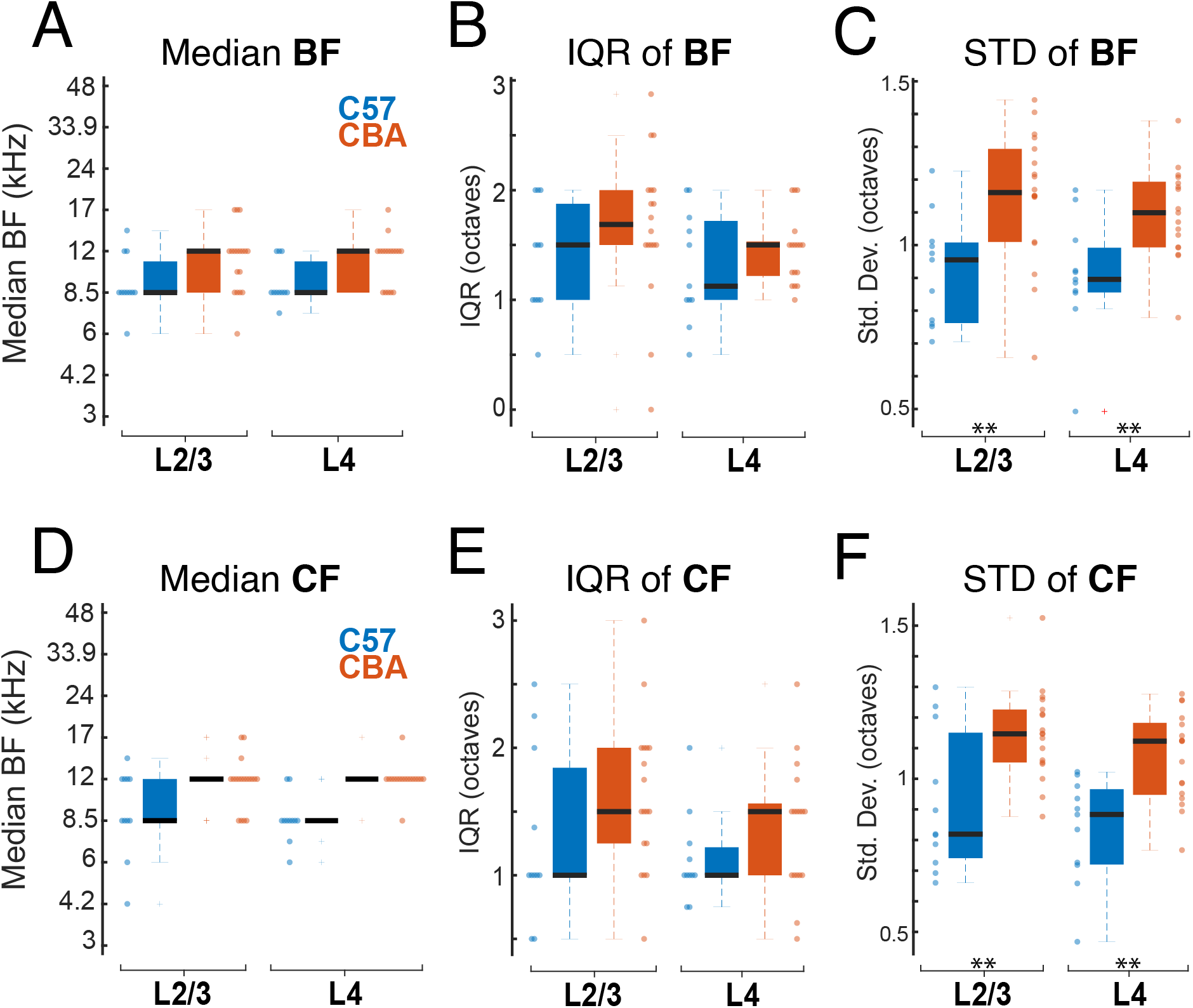
Tuning heterogeneity is higher in CBA mice. **A)** Median BF for each FOV in L2/3 (left) and L4 (right). Each dot represents one FOV. Blue indicates C57 mice and orange indicates CBA mice. **B)** IQR of BF in each FOV for L2/3 (left) and L4 (right). **C)** IQR of BF in each FOV for L2/3 (left) and L4 (right) populations. ** indicates significant difference at p<0.01 (Wilcoxon rank sum test or two-same t-test depending on normality). **D,E,F)** Conventions as in **A,B,C** but for CF.

While CF and BF are generally well correlated, we demonstrated (Fig. 2B) that they often differ due to FRA nonlinearities. We thus also quantified the median, IQR, and STD of CFs from each FOV. We found precisely the same trends between mouse strains as observed with BF. Specifically, we find that median CF was generally the same in both layers of C57 mice (8.5 kHz) and across both layers of CBA mice (12 kHz) (Fig. 3D). IQR_CF_ was higher in CBA mice that in C57 mice in both L2/3 and L4 (Fig. 3E). STD_CF_ was significantly higher in CBA mice than in C57 mice in both L2/3 and L4 (Fig. 3F) (p_L2/3_<0.004, p_L4_<10^−3^, two-sample t-test). These results reinforce our finding that frequency tuning heterogeneity of neuronal populations is present in both strains of mice, yet larger in CBA mice. The larger spread of preferred frequencies in CBA mice compared to C57 mice is likely due to the deficit of high frequency tuned neurons observed in neuronal populations of C57 mice.

### Increased tuning heterogeneity in CBA mice is present at multiple length scales

We next investigated whether the differences in tuning heterogeneity between strains is restricted to small length scales or is maintained over larger populations of neurons. We calculated IQR_BF_ and IQR_CF_ at several different length scales, where the IQR is computed in a neighborhood surrounding each individual neuron. One neuron was picked as the central neuron and then IQR was computed from BFs of neurons within a radius ranging from 50 to 400 µm around that neuron (Fig. 4A). This process was repeated for each neuron in the FOV, using each one as the center of the local neighborhood. We found that CBA mice maintain a higher extent of frequency tuning heterogeneity across all length scales up to the size of our FOVs than C57 mice (Fig. 4B). Furthermore, we observed slightly lower IQR_BF_ values at smaller length scales, particularly in L2/3, indicating a small-scale localization of similarly tuned neurons.

**Figure 4.**
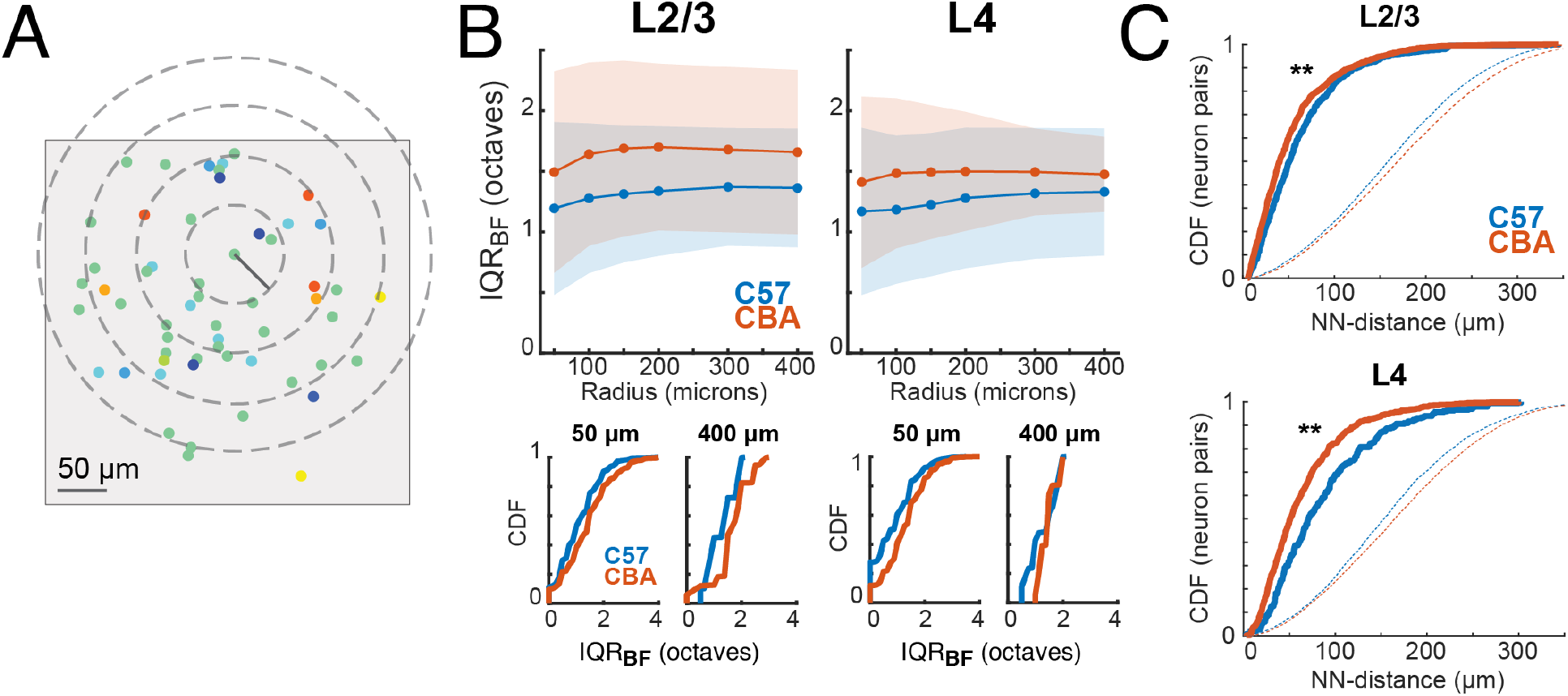
CBA mice have more tuning heterogeneity across length scales. **A)** Schematic illustrating concentric circles (“neighborhoods”) of varying radii around a single neuron. Gray box indicates FOV. 300µm and 400µm circle not shown. **B)** *Top*, Mean IQR_BF_ for each mouse strain at each length scale. Shading represents STD. *Bottom*, CDFs of IQR_BF_ for each mouse strain at short (50µm) and long (400µm) length scales. **C)** CDFs of same-tuned nearest-neighbor distances for neuron pairs in each mouse strain. Dashed lines indicate randomly sampled pairwise distances from the neuronal populations. ** indicates significant difference between strains at p<0.01 (Wilcoxon rank sum test).

We further analyzed the spatial heterogeneity of tuning preference by performing analysis outlined in (Zeng et al., 2019). We first computed the nearest-neighbor distance for each neuron in a FOV by finding the minimum distance to another neuron with the same tuning. We compared these distributions to random distributions where an equal number of pairwise distances were randomly sampled from the possible pairwise distances between tuned neurons. We found that both CBA and C57 mice had nearest neighbor distances significantly smaller than random. CBA mice had smaller nearest-neighbor distances than C57 mice (Fig. 4C) which serves as further support that despite the increased heterogeneity in CBA FOVs, the similarly tuned cells are more densely localized. Together these data show that the spatial heterogeneity in frequency preference does not depend on mouse strain and thus is a feature of rodent auditory cortex.

### Signal correlations are higher in C57 mice

We next investigated if the slight differences we observed in receptive fields of each mouse strain were reflected in measures of functional connectivity. We calculated signal correlations which infer similar feedforward input to neuronal pairs by looking at the covariance between receptive fields of each pair. We observed non-normal distributions with a bias towards positive values for signal correlations. We found that signal correlations (SC) are higher in C57 mice in both L2/3 and L4 neuronal populations (Fig. 5A) (L2/3 mean: SC_C57_=0.17, SC_CBA_=0.14; L4 mean: SC_C57_=0.16, SC_CBA_=0.10). This indicates that overall, neuronal pairs in C57 mice have more similar inputs than in CBA mice. Next, we calculated noise correlations (NC) which infer direct connections between neurons or shared sources of activity perturbations. NCs centered closer to zero than SCs with a slight bias towards positive values. NCs CBA mice trended slightly higher in L2/3 than in C57 mice (Fig. 5B) (L2/3 mean: NC_C57_=0.04, NC_CBA_=0.06; L4 mean: NC_C57_=0.04, NC_CBA_=0.03).

**Figure 5.**
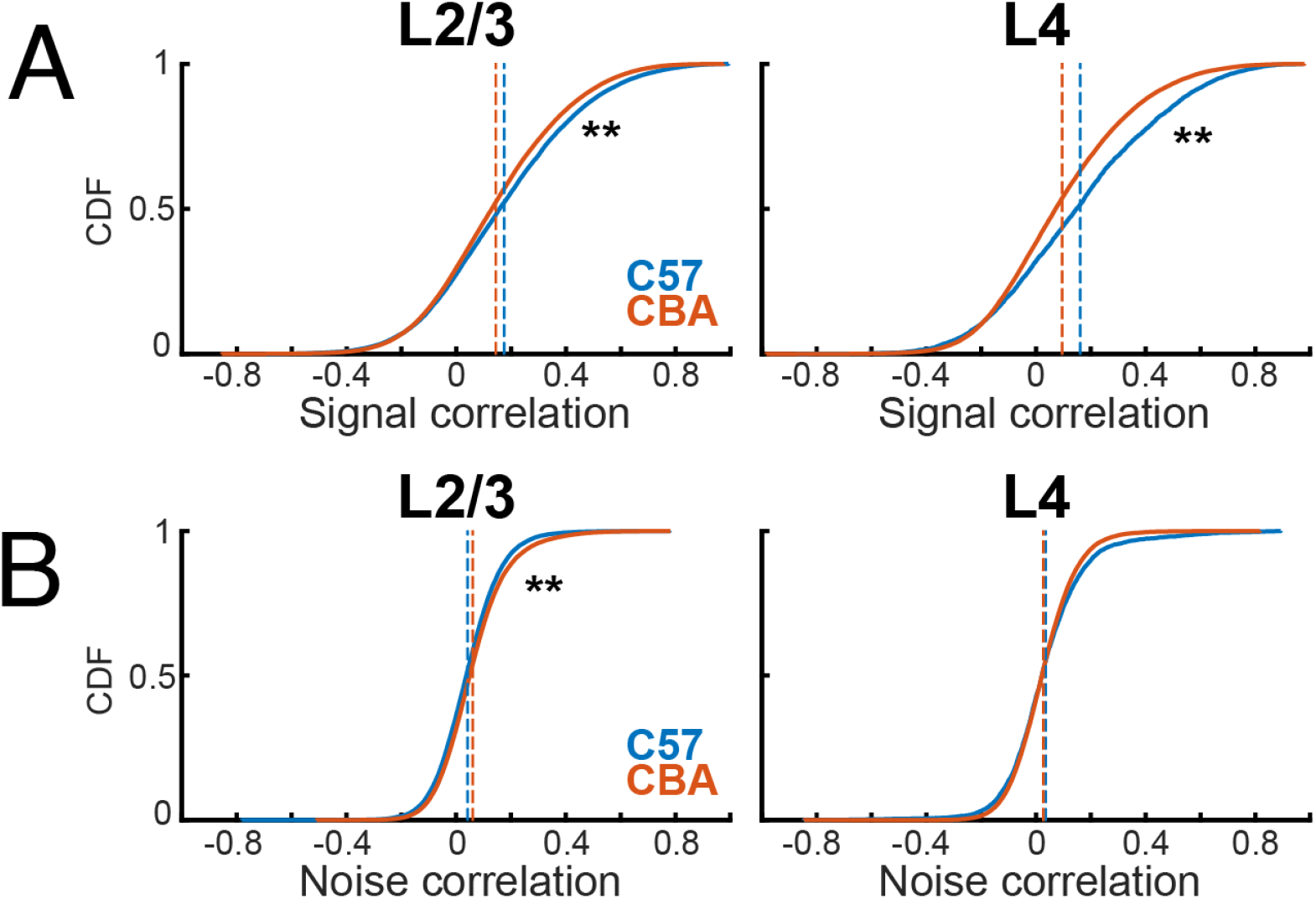
Signal and noise correlations across strains. **A)** CDFs of signal correlations in both strains in L2/3 (*left*) and L4 (*right*). Dashed lines indicate distribution mean. ** indicates significant difference at p<0.01 (Wilcoxon rank sum test). **B)** Conventions as in **A**, but for noise correlations.

Since C57 mice have a decreased frequency representation of high frequencies we next investigated if changes in pairwise correlations were similar for cell pairs across the hearing range. We thus separately calculated pairwise correlations between cells with frequency preference in different octave bands. SCs in L2/3 were similar in 3-6KHz and 6-12kHz band between genotypes but were higher in C57 mice in 12-24 and 24-48kHz (Fig. 6A). In L4, consistent with our overall observations, SCs were higher in most frequency bands, except for the 24-48kHz band were only a trend was observed (Fig. 6B). These results show that C57 mice have higher SCs than CBA mice across most of the hearing range. NCs were larger for frequency bands <24Khz in CBA mice than in C57 mice (Fig. 6C), but NCs were largely similar in all frequency across the hearing range in L4 (Fig. 6D). Together, these results indicate that while overall pairwise correlations between CBA and C57 are similar small differences exist and at that these differences are present in the middle frequency ranges which do not show reduced representation in C57. These results suggest that there are subtle intrinsic differences in A1 processing between mouse strains.

**Figure 6.**
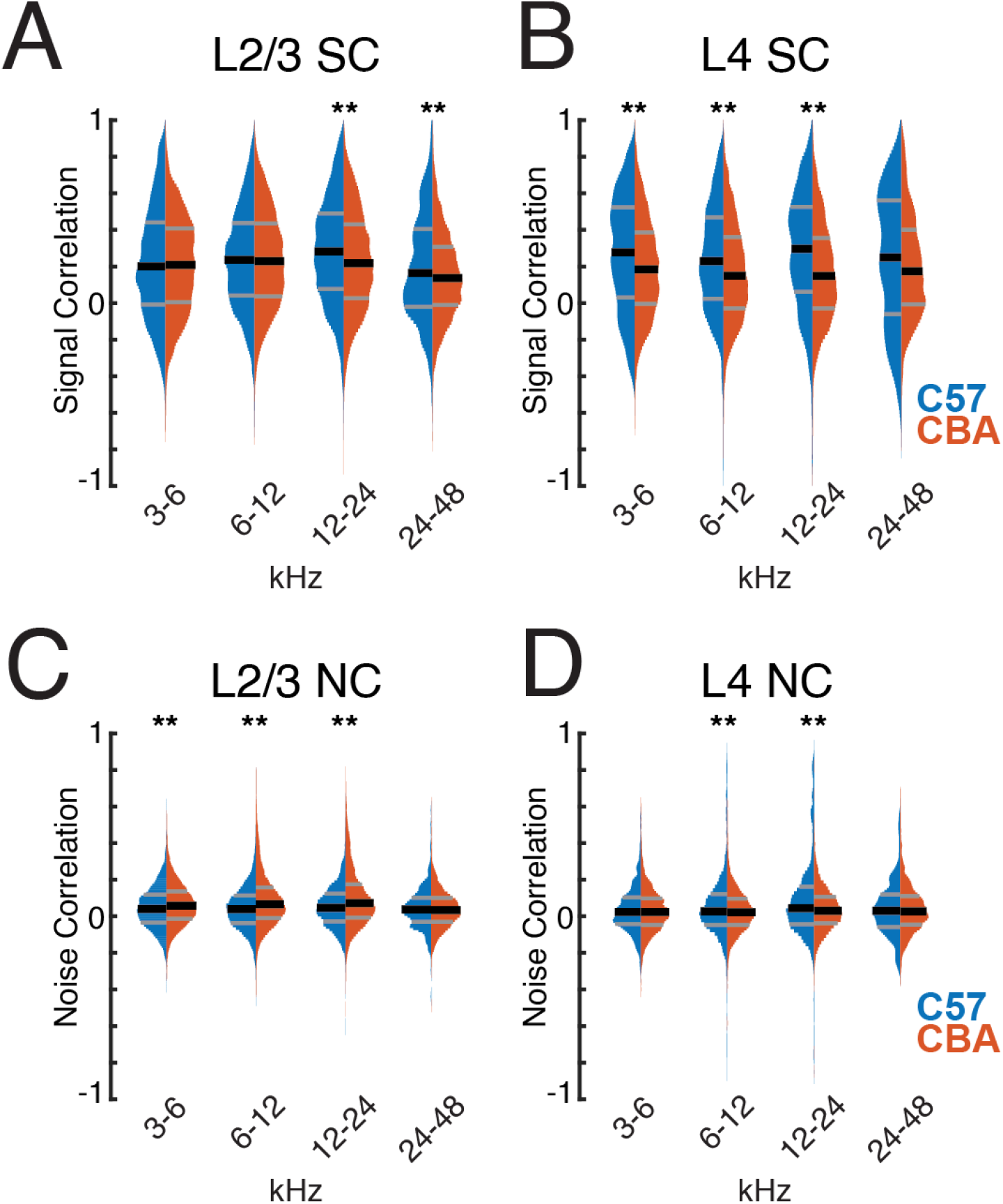
Signal and noise correlations of similarly tuned neurons. **A)** Signal correlation distributions of similarly tuned neurons in L2/3 of both strains. ** indicates significant difference at p<0.01 (Wilcoxon rank sum test). **B)** Conventions as in **A**, but for L4. **C)** Noise correlation distributions of similarly tuned neurons in L2/3 of both strains. ** indicates significant difference at p<0.01 (Wilcoxon rank sum test). **D)** Conventions as in **C**, but for L4.

Given the observed tuning diversity in ACX neuronal populations, we next investigated how SCs differ depending on the BF relationships of the cell pair. We quantified the SCs of cell pairs as a function of difference in preferred frequency. We found that in L2/3 populations of both strains SCs drop off as BFs differ between neurons (Fig. 7A), even if the difference is as small as 0.5 octaves, and then remain relatively constant with respect to ΔBF. In L4 populations of both strains, SCs again drop off at a ΔBF of 0.5 octaves (Fig. 7B) and then remain relatively constant at differences greater than 1 octave. In addition to the slight shifts in means, both L2/3 and L4 populations of both strains have a wide distribution of SCs at all ranges of ΔBF. These results indicate that despite differences in receptive fields and tuning diversity of both mouse strains, pairwise activity relationships in activity seems largely unchanged.

**Figure 7.**
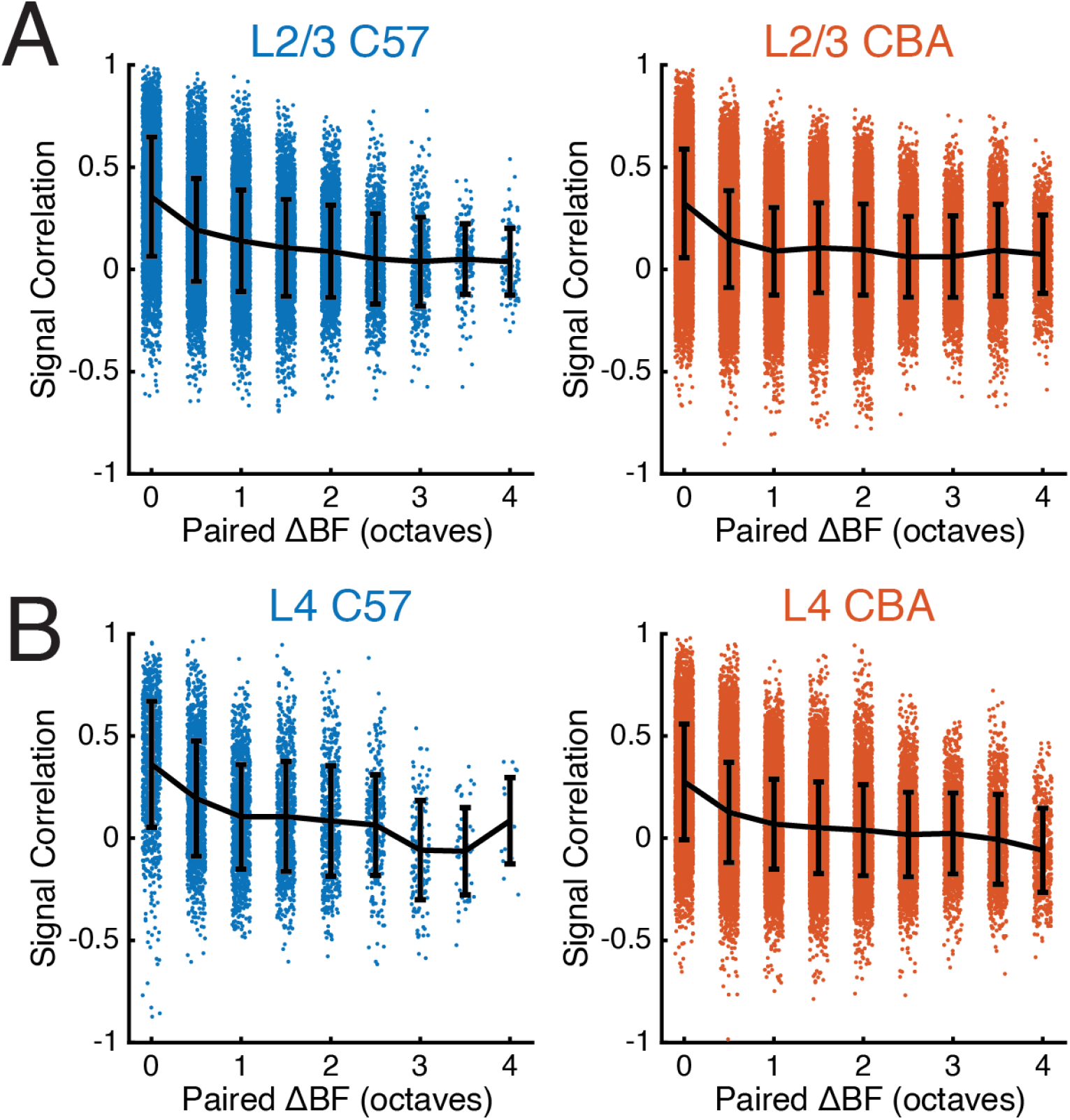
Signal correlations decrease as tuning preference deviates. **A)** Signal correlation distributions of neuron pairs with respect to their difference in preferred frequency in L2/3 of both strains. **B)** Conventions as in **A**, but for L4.

## Discussion

Using in vivo 2-photon imaging we show that A1 in both the C57 and CBA strain of mice contains neurons largely similarly tuned and responsive for sound frequencies but that CBA mice contain more neurons selective for high frequencies. Moreover, we show that the spatial representation of sound frequency representation in A1 is heterogeneous in both strains of mice. Thus, the observed heterogeneity in prior studies using C57 mice is not due to changes in the cochlea and subsequent central plasticity in these mice and indicates that heterogeneous representation of sound preference is a feature of rodent auditory cortex.

C57 mice show progressive degeneration of the basal turn of the cochlea that becomes evident on a histological level by 3 months (Park et al., 2010). In addition, auditory brainstem response (ABR) differences can be observed at ~10 weeks (Ison et al., 2007; Martin del Campo et al., 2012). However, both measures are relatively coarse, thus it is likely that subtle functional deficits are present at even earlier ages. The mice in our study were within this range consistent with their relative normal hearing. The main deficit we observed was a relative paucity of high-frequency (>32kHz) responding cells in C57 mice consistent with an emerging degeneration of peripheral high-frequency hearing and the changes on the ABR level (Ison et al., 2007; Martin del Campo et al., 2012).

Functional and molecular differences between strains are present on the brainstem level at even younger ages but some of these could reflect strain differences unrelated to hearing loss. For example cells in the VNTB fire at lower rates in C57 mice than CBA mice at young ages (Sinclair et al., 2017) and deficits in the efferent feedback precede overt hearing deficits and could be observed by 8 weeks (Frisina et al., 2007; Zhu et al., 2007). By 6 months of age C57 mice also show a loss of parvalbumin immunoreactivity in A1 and AAF suggesting a loss or hypofunction of fast spiking (PV) inhibitory interneurons (Martin del Campo et al., 2012; Brewton et al., 2016). However, C57 mice show a higher number of PV cells than CBA mice at young ages indicating baseline strain differences in cortical circuits (Brewton et al., 2016). A loss of inhibition with age would be expected to result in changes in the functional responses, such as a broadening of FRAs. While we did observe a bandwidth difference between strains, our data showed higher frequency selectivity in C57 mice than CBA mice consistent with the higher baseline number of PV cells in these mice (Brewton et al., 2016). Additionally, any decrease in PV immunoreactivity in C57 mice might have not impaired the function of PV cells, or decreased PV cell function in C57 mice might have been compensated for by other classes of interneurons.

We observed both onset and offset responses consistent with prior reports (Liu et al., 2019). Possible due to our usage of shorter tone stimuli or SAM tones we observed a lower fraction of offset responsive neurons than our prior study (Liu et al., 2019). Furthermore, here we used a stricter criterion for off-responsive neurons such that on-responsive neurons were disqualified from also being off-responsive neurons. Nevertheless, we observed an altered ratio of onset to offset neurons with C57 mice having fewer offset neurons that CBA mice. While onset and offset responsiveness is determined by distinct thalamocortical circuits, inhibitory circuits are thought to play a role in enhancing offset responses (Liu et al., 2019). Thus, our observation here could indicate hypofunction of inhibition. Given that we do not observe broadening of receptive fields in C57 mice as compared to CBA mice, temporal effects of inhibitory hypofunction reflected in offset responses might be more sensitive indicator of inhibitory hypofunction than pure tone bandwidth.

Thus, consistent with prior anatomical and ABR data our single cell results suggest that at ~10 weeks of age there is no large effect of hearing loss on A1 of C57 mice besides the paucity of high frequency neurons in mid-frequency regions. However, it is possible that neuroplastic changes have already been taking place to adjust the tuning of these former high-frequency neurons, compensating for peripheral changes.

Our results show that a local diversity of frequency tuning is present in mice of both strains. This confirms observations using in vivo 2-photon Ca^2+^-imaging under various conditions in multiple labs. Initial studies using synthetic dyes in anesthetized animals described a local heterogeneous frequency preference (Bandyopadhyay et al., 2010; Rothschild et al., 2010) in L2/3. These results in L2/3 have been confirmed using the sensitive synthetic dye Cal-520 (Li et al., 2017; Tischbirek et al., 2019; Zeng et al., 2019) in anesthetized and awake mice and rats, with the ultrasensitive genetically encoded calcium indicator (GECIs) GCaMP6s in awake mice (Meng et al., 2017; Liu et al., 2019), as well as by in vivo patch clamp studies (Maor et al., 2016). In addition to primary auditory cortex, L2/3 of ACX areas such as the anterior auditory field and secondary ACX were also shown to contain fractured tonotopic organization (Liu et al., 2019). Interestingly, using low sensitivity indicators (GCaMP3) revealed a high degree of local similarity (Issa et al., 2014) suggesting that strongly responding neurons in a local field might be very similarly tuned possibly due to subsampling certain classes of L2/3 neurons (Meng et al., 2017). Investigation of the tuning of MGB afferents in A1 showed that the local frequency distribution of MGB terminals was very heterogeneous (Vasquez-Lopez et al., 2017; Liu et al., 2019). Thus at least in mouse a large degree of local tuning diversity is present on multiple levels in the auditory cortex. Further studies using the synthetic dye Fluo-4 revealed that L4 showed less local tuning diversity than L2/3 (Winkowski and Kanold, 2013) consistent with electrophysiological results (Guo et al., 2012; Kanold et al., 2014). The increased diversity in L2/3 over L4 suggest that processing complexity might increase in superficial layers consistent with recent findings in humans (Moerel et al., 2018; Moerel et al., 2019). Interestingly, studies in marmoset find local tuning homogeneity in marmoset A1 which indicates species differences in local A1 organization (Zeng et al., 2019).

In general, the degree of heterogeneity in the local frequency organization varies between prior rodent studies. In particular, recent imaging using the synthetic dye Cal-520 found no difference between layers (Tischbirek et al., 2019). While these differences between studies might be due to differences in the experimental conditions, cell inclusion criteria, range of tonal stimuli and sound levels used, other factors might play a role. We speculate that the differences in local frequency preference might reflect plastic changes due to different rearing environments. Exposure to sounds in development can alter the functional organization of A1 (Zhang et al., 2001, 2002; Chang and Merzenich, 2003). Since rodent housing can be loud and might vary considerably between institutions (Lauer et al., 2009), it is possible that differences seen between studies reflect sensory adaptations to the local environment or low levels of hearing loss induced by personal activity in the local environment. Support for this hypothesis comes from studies in other systems. Local heterogeneity of tuning in L2/3 is a feature of not only rodent A1 but also of primary visual and somatosensory cortices (Ohki and Reid, 2007; Bonin et al., 2011; LeMessurier et al., 2019). For example, S1 local L4 neurons show very similar tuning, yet L2/3 neurons are more diverse (LeMessurier et al., 2019). This difference between layers was dependent on mice living in impoverished versus enriched environments suggesting that experience is shaping the differential representation of stimulus properties in L4 and L2/3. Our study investigated A1 in two strains of mice raised in the same sound environment. Our colony room is equipped with silent door latches and sound dampening panels reducing the occurrence of loud transient and steady state background sounds (Lauer et al., 2009). Our results in mice of two strains using genetically encoded Ca^2+^ indicators show similar functional organization of A1 as our prior studies using synthetic dyes (Winkowski and Kanold, 2013). Thus, the local tuning diversity especially in L2/3 is a common property of rodent auditory cortex.

Our measurements of signal correlations were generally less positive than previously reported results (Winkowski and Kanold, 2013). In contrast to these prior studies we calculated the covariance of responses across multiple sound levels rather than one sound level in order to more fully capture the interactions between neurons. Receptive fields of ACX neurons are highly nonlinear with respect to sound level, and as a result our inputs to the covariance measure are more variable than when using one sound level.

In summary, our results suggest that A1 of C57BL/6 in adult mice is very similarly responsive and similarly organized to A1 of CBAxC57 F1 mice. Thus, while aspects of cellular tuning differ between strains, the differences in cortical processing at this age is modest at best and largely restricted to a paucity of high frequency neurons.

## Acknowledgements & Contributions

ZB, DEW, and POK designed research. DEW performed experiments. ZB analyzed the data. ZB, DEW, and POK discussed the results. ZB and POK wrote the manuscript. Supported by NIH U01NS090569 (POK), U19NS107464 (POK), RO1DC009607 (POK), DOD W81XWH-16-1-0143 (POK, DEW).

